# Chromosomal markerless integration of anthelmintic Cry proteins into the *Bacillus thuringiensis* genome

**DOI:** 10.64898/2026.04.24.720002

**Authors:** Kelly A. Flanagan, Nicholas Cazeault, Hanchen Li, Elizabeth Kass, Katherine L. Petersson, Gary R Ostroff, Raffi V. Aroian

**Affiliations:** Program in Molecular Medicine, UMass Chan Medical School, Worcester, MA 01605; Department of Fisheries, Animal, and Veterinary Sciences, University of Rhode Island, Kingston, RI, 02881

**Keywords:** *Bacillus thuringiensis*, markerless integration, allelic exchange, Cry proteins, anthelmintics, IBaCC, nematicidal bioactivity

## Abstract

*Bacillus thuringiensis* (Bt) is a Gram-positive bacterium that during sporulation produces insecticidal Crystal (Cry) proteins, which play a major role in insect control today. Some Bt Cry proteins, e.g., Cry5Ba, target nematodes and, when given orally, can cure animals of gastrointestinal nematode (GIN) parasites. To eliminate concerns about treating humans and animals with spores and live bacteria, we developed an asporogenous system for scalable and safe Cry protein delivery called IBaCC (Inactivated Bacteria with Cytosolic Crystal(s)), which results in production of a bioactive crystal and a dead bacterium. However, to date, IBaCC involves expression of Cry proteins from antibiotic-selectable plasmids to ensure maintenance. Here, we develop and validate tools for markerless and stable integration and expression of Cry proteins in Bt. We markerlessly integrate an expression construct for Cry5Ba into either the *spo0A* or the *sigK* locus and demonstrate robust Cry5Ba expression. We also integrate our Cry5Ba expression construct into both loci simultaneously, increasing expression further. We demonstrate that an expression construct for a second anthelmintic Cry protein, Cry21Aa, can be integrated either alone or in combination with Cry5Ba in a single Bt strain. We furthermore show that these markerless integrants are stable in the absence of a selectable marker. These integrated strains, processed to IBaCC, demonstrate excellent ex vivo nematicidal bioactivity toward the larval stages of the sheep GIN parasite *Haemonchus contortus* and adult stages of the human hookworm GIN parasite *Ancylostoma ceylanicum*. This study demonstrates the successful markerless integration of 1-2 identical or dissimilar Cry proteins into Bt. These Cry integrants, in which genes essential to sporulation are deleted or replaced, provide robust Cry expression, stability, and bioactivity. These studies represent an important advance in Bt genetics and toward a safe, deployable, and cost-effective anthelmintic therapy to treat GIN parasitic infections in humans and animals.

## Introduction

The gram-positive spore-forming soil bacterium *Bacillus thuringiensis* (Bt) is the dominant and most important biologically produced insecticide, extensively used in agriculture and vector control today (Rakesh, Kalia, and Ghosh 2023; Gassmann and Reisig 2023; Azizoglu et al. 2023). The main insecticidal factors used to kill insects in Bt sprays and in transgenic plants are crystal, or Cry, proteins that are often produced during sporulation and are so named because they form large parasporal crystals (Deng et al. 2014; Sanahuja et al. 2011). Bt and its Cry proteins have a strong safety record for mammalian safety *per os* (Raymond and Federici 2017; Oliveira-Filho and Grisolia 2022). More than 800 insecticidal Cry proteins categorized into > 50 families have been studied, and Bt has been deployed against insect pests for more than six decades (Nester et al. 2020; Crickmore et al. 2021).

Some Cry proteins, *e.g.*, Cry5Ba, have been shown to specifically target nematodes (Marroquin et al. 2000; Wei et al. 2003), a discovery that opened the possibility of using Cry proteins against nematode pests. Nematodes, namely soil-transmitted helminths or STH and gastrointestinal nematodes or GINs, are amongst the most important pathogenic and common parasites of humans and animals, leading to significant morbidity and in some cases mortality (Khurana, Singh, and Mewara 2021; WHO 2023; Jones and Garcia 2019). Deworming (anthelmintic) drugs have incomplete efficacies and are prone to resistance development by GIN parasites, mandating the search for new anthelmintics (Ahuir-Baraja et al. 2021; Tinkler 2020). Oral dosing of Cry5Ba spore-crystal lysates and purified protein has been shown to cure human and animal parasitic nematode infections in rodents, dogs, and pigs, indicating that Bt Cry proteins could help solve the unmet need for new anthelmintics (Hu et al. 2018; Urban et al. 2013; Cappello et al. 2006; Hu et al. 2012).

To develop Cry proteins as oral therapeutics, we recently published the development of IBaCC (for *I*nactivated *Ba*cteria with *C*ytosolic *C*rystal(s)) in which a nonsporulating Bt strain expresses Cry proteins during vegetative growth and is subsequently inactivated (killed) using food-grade terpenes. IBaCC, a dead bacterium with a live Cry protein payload, is an industrially scalable production system that allows Cry proteins to be produced cheaply and simply and then delivered orally without the necessity of using live bacteria and all the associated disadvantages (Li et al. 2021; Chicca et al. 2022; Urban et al. 2021; Hoang et al. 2024). IBaCC containing Cry5Ba, Cry14Ab, and Cry14Ac crystals has been shown to have significant therapeutic if not curative effects on one or more parasitic infections by hookworm, ascarids, whipworm, and/or *Haemonchus* in vertebrate hosts that include rodents, pigs, sheep, and horses (Chicca et al. 2022; Li et al. 2021; Sanders et al. 2020; Urban et al. 2021; Hoang et al. 2024).

For all anthelmintic and IBaCC Cry protein studies involving recombinant strains to date, Cry protein expression has been driven from multi-copy extrachromosomal plasmids that require antibiotic resistance selection for maintenance. For large-scale production of an IBaCC therapeutic, such plasmids are not ideal as they raise cost (antibiotic use), require removal of residual antibiotics, potentially spread antibiotic resistance genes, and complicate regulatory approval (Arsene et al. 2022; Mignon, Sodoyer, and Werle 2015; Jian et al. 2021; Vidal et al. 2008). We therefore set out to establish for the first time markerless integration in Bt. We explored the utility of integrating Cry genes singly or in combination into the Bt genome without leaving behind antibiotic selectable markers. We further ascertain whether such integrants express Cry proteins well, whether they are stable in the absence of selection, and whether the Cry proteins produced are bioactive against parasitic nematodes. These results have important implications for the development of Cry-protein anthelmintics, next-generation insecticides, and Bt biology.

## Materials and Methods

### General microbiology

*Escherichia coli* 5α strains (New England Biolabs, NEB), each harboring a specific plasmid, were used for cloning and maintained with Luria-Bertani (LB) medium including ampicillin (100 μg/ml) at 37° C or erythromycin (350 μg/ml) at 30° C. *E. coli dam^-^/dcm^-^* (NEB) cells were used to generate nonmethylated plasmids for Bt transformations and similarly maintained on LB medium with antibiotics when appropriate. Acrystalline, *spo*^+^ Bt strains, each cured of large *cry*-encoding plasmids, were each obtained from the Bacillus Genome Stock Center (Columbus, OH, USA; see Table S1). All Bt strains were maintained at 30° C with LB medium, including erythromycin (10 μg/ml) for plasmid maintenance when appropriate. Solid media contained 1.6% agar. Optical densities (OD_600_) were measured in cuvettes with a SmartSpec 3000 spectrophotometer (Bio-Rad, CA) and reported as an average of technical duplicates at each time point.

### Phase-contrast microscopy

An Olympus BX60 microscope equipped with a UPlanFl 100x/1.30 Oil Ph3 objective was used to both confirm *spo* phenotypes of Bt strains grown on solid media >48h (Figures 1 and 2) and to visualize recombinant Cry crystals in *cry*-integrated strains (Figures 2 and 6). Images were captured using μManager (Edelstein et al. 2014) software and further visualized with Fiji (Schindelin et al. 2012).

**Figure 1.**
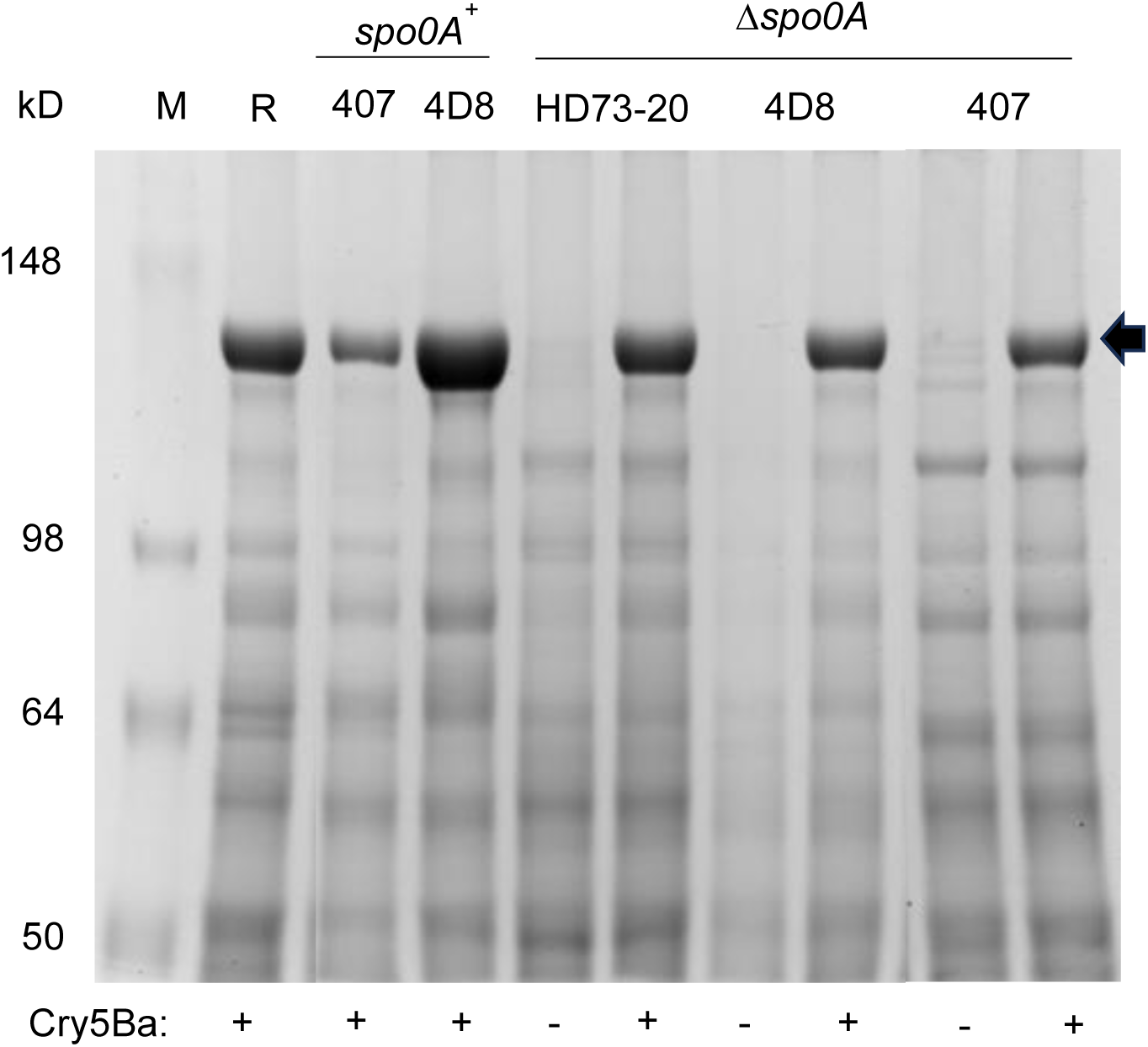
Comparison of *spo^+^* and *spo^-^* Bt strains for Cry5Ba content. SDS-PAGE analysis of Cry5Ba production in *spo0A^+^* and *spo0A^-^* Bt strains after 48h of culture in LB. Expression of Cry5Ba from plasmid pHY159 in *spo^+^* cells (*spo0A^+^*) is compared to expression of Cry5Ba from plasmid pHY159 in which the *spo0A* gene has been deleted (Δ*spo0A*). Cry5Ba - = pHT3101 vector with no insert; Cry5Ba+ = pHY159, Cry5Ba expression plasmid. M = protein marker, R = reference strain, 407 Δ*spo0A*::*kan*.

**Figure 2.**
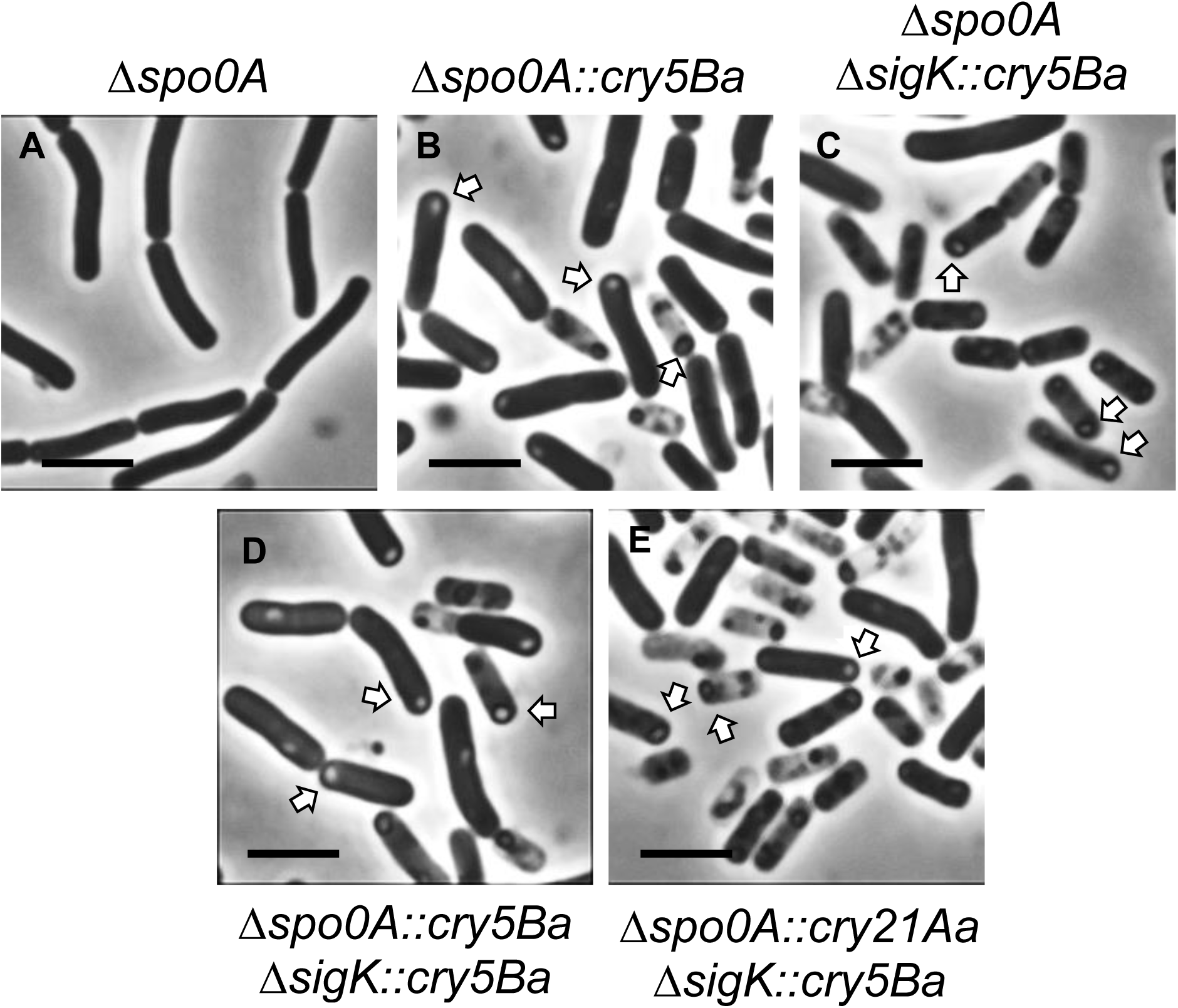
Phase-contrast microscopy of markerless, asporagenous *Bt* 407 strains with *cry* genes integrated in the chromosome. A) Stationary cultures of Δ*spo0A* cells without a *cry5Ba* insert cannot sporulate and are void of Cry^+^ inclusion bodies. B-D. *spo^-^* cells with only *cry5ba* inserted into the chromosome to replace *spo0A* (B), or *sigK* (C), or both *spo0A* and *sigK* (D). (E) *cry5Ba* and *cry21Aa* replacing the *sigK* and *spo0A* loci respectively showing a prominent crysta*l.* Arrows point to crystals (inclusion bodies). Crystals in Δ*spo0A* cells with *cry21Aa* integrated alone are not visible by microscopy at this scale. Scale bar = 5 µm.

### Plasmid construction

Phusion polymerase (NEB) was used for PCR amplification of plasmid inserts, and HiFi DNA Assembly master Mix (NEB) was used for plasmid assembly in all cases (all primers are given in Table S2). To construct the Cry expression plasmids for Cry5Ba and Cry21Aa, published plasmids were used as template for PCR amplification (Marroquin et al. 2000; Wei et al. 2003). Plasmid pHY159 harbors an approximately 4.9kb expression construct for Cry5Ba; this includes the promoter region of *cry3A* (P*_cry3A_*) fused to the coding sequence of *cry5Ba* and the 3’ untranslated region of *cry5Ba* (Li et al., 2021). Since this original construct contains 6 amino acids from Cry3Aa, we also generated a construct in which the *cry3A* promoter region (-635 to +0) was fused to each PCR-amplified coding sequence starting at nucleotide 1 of *cry5Ba* or *cry21Aa,* then fused to the *cry5Ba* terminator. These expression constructs were inserted into pHT3101 (Li et al. 2021; Lereclus et al. 1989) to clone for transformation into Bt and respectively are called P*_cry3A_-cry5Ba* or P*_cry3A_-cry21Aa* (see Table S3).

We constructed *spo0A* deletion plasmids as follows. Generally, ∼ 2 kb strain-specific Δ*spo0A* loci were each cloned into the integration plasmid pHY304 (kindly provided by Dr. Craig Rubens’ laboratory, Seattle Children’s Research Institute) (Jones, Needham, and Rubens 2003; Pflughoeft, Sumby, and Koehler 2011), transformed into the appropriate Bt recipient and introduced into the chromosome by routine allelic exchange. pHY304 harbors a temperature-sensitive origin of replication and has been successfully utilized for allelic exchange in other *Bacillus* species (Corsi et al. 2020; Yim and Rubens 1998). Chromosomal DNA was first isolated (Promega Wizard) from each acrystalline Bt strain to use as PCR template. Primers (see Table S2) were designed to the genomes of Bt 407 (Accession: NC_018877.1), HD73-20 (Accession: CP004069.1), and HD-1 (Accession: CP158703.1) to amplify ∼1kb of the 5’- and 3’ arms flanking the *spo0A* sequences with overlapping primers. When assembled, the PCRs overlap, deleting *spo0A* and its 5’ ribosomal binding site. Specifically, the 804 bp sequence deleted from the newly assembled Δ*spo0A* locus includes the last three nt in the -10 binding site of its essential σ^A^ promoter through the first 775bp of the 831bp gene (-29 through 775 of the coding sequence), leaving behind a defective promoter and 3’ gene fragment of 56 bases without a functional start site for transcription or translation.

We performed integrations of *cry5B* or *cry21Aa* genes into one of two loci, *spo0A* or *sigK*. As with spo0A above, 5’ and 3’ arms flanking *sigK* were PCR amplified with overlapping primers (see Table S2) to assemble together, deleting *sigK*, and subcloned into pBluescript. ∼ 2kb null deletion (Δ) loci for *spo0A* and *sigK* were each subcloned to increase targeting accuracy of homologous recombination. A unique EcoRV restriction enzyme site was introduced into these overlapping primers for the subsequent insertion of *cry* genes into Δ*spo0A* or Δ*sigK* pBluescript subclones (see Table S3 for pHY306 and pHY311 respectively). The P*_cry3A_-cry5Ba* or P*_cry3A_-cry21Aa* expression construct, including the *cry3A* promoter and *cry5Ba* terminator, was amplified with primers prKF150 and prKF151 (see Table S2) from pHY300 or pHY301, respectively, and inserted into each of these EcoRV-digested pBluescript derivatives to generate subclones of Cry expression constructs at each deleted locus. From these subclones, entire replacement loci (*Δspo0A*::*cry5Ba*, Δ*spo0A*::*cry21Aa*, Δ*sigK*::*cry5Ba*) were PCR-amplified (see Table S2 for pHY304-specific primers) and finally assembled into BamHI-and HindIII-digested pHY304. *E. coli* 5α cells were transformed with DNA Assembly reactions and selected on erythromycin. All inserts from start to end were confirmed by Sanger sequencing. DNA sequence analyses were performed using Geneious Prime software (www.geneious.com). All plasmids for expression and integration used in this study are listed in Table S3.

### Bt transformation and integration

Electroporation-competent Bt cells were prepared and transformed using published methods (Lereclus et al. 1989; Macaluso and Mettus 1991; “Bacillus Genetic Stock Center Catalog of Strains” 1999). Briefly, electrocompetent Bt were transformed with 1-2 μg nonmethylated plasmids (see Table S3) using with the Electro Cell Manipulator 630 (Harvard Apparatus, BTX) set at 2500 V with 25 μF capacitance. Transformations of all Bt strains with control or Cry expression plasmids were selected on erythromycin (see Table S4) and then confirmed by PCR verification of each plasmid.

To induce allelic exchange with pHY304-derivatives in transformed strains, erythromycin-resistant transformants were subjected to an overnight temperature shift, as previously described (Jones, Needham, and Rubens 2003) with slight modifications. In brief, a single erythromycin-resistant isolate of each transformation was grown in 5ml liquid LB with erythromycin at 41 °C (Pflughoeft, Sumby, and Koehler 2011), which was then streaked onto LB erythromycin agar plates at 41 °C to acquire isolated colonies with integrated plasmid. Next, a single isolate from this plate was cultured in LB without erythromycin at 30 °C overnight, and dilutions were plated onto solid LB without erythromycin. For each new integrated Bt strain, up to 50 colonies were patched onto separate solid LB plates, with and without erythromycin to identify erythromycin-sensitive isolates (integrants). Exhausted colonies, sampled from the center, and liquid cultures at the end of incubations were assessed alongside *spo^+^* original strains for evidence of mature spores. The *de novo* loss of sporulation in Δ*spo0A* and Δ*sigK* deletions and integrations were confirmed for their inability to sporulate (*spo*^-^) by the lack of phase-bright spores by phase-contrast microscopy. These isolates were confirmed by PCR of the *spo0A* and/or *sigK* locus and validated via sequencing. All Bt strains constructed in this study are listed in Table S4.

### Protein expression and preparation

All expression cultures were incubated at 25° or 30° with 125 or 250 rpm shaking agitation, as indicated. For Cry protein expression cultures, 3X LB medium, which contained erythromycin at 10 ug/mL when appropriate, was inoculated with Bt and cultured at 30° 250rpm for 2-3 days, as indicated. Small-scale cultures with 10ml medium were grown in 25mm-diameter culture tubes. For greater aeration and rapid growth, larger cultures were instead grown in beveled flasks at 10% flask volume (*e.g.* 25ml in 250ml flasks). Inoculations were typically done with a 1:100 dilution of overnight cultures. IBaCC preparations were generated as described (Li et al. 2021) and quantified by SDS-PAGE using bovine serum albumin (BSA) standards. Cell and BSA samples were denatured in 1X Laemmli sample buffer (Alfa Aesar) at 95 °C 5-10 minutes, and denaturing SDS-PAGE was performed using the

Mini-Protean system (Bio-Rad), including 7.5% acrylamide hand-cast and precast TGX gels (Bio-Rad). All protein gels were stained with Coomassie based solutions (Bio-Rad SafeStain or IBI Scientific Quick Protein Stain) and imaged with the Ingenius3 gel documentation system (Syngene). Densitometry for protein quantification was performed using Fiji (ImageJ) software. Negative control Bt preparations that lacked Cry proteins were used for background subtraction.

### Mass Spectrometry

50ug and 100ug total protein samples, determined by densitometry of gel-resolved proteins relative to BSA, from strain HY-BT218 (Δ*spo0A*::*cry21Aa* Δ*sigK*::*cry5Ba*, see Table S4) were applied to separate 5-well, 8% acrylamide gels and electrophoresed at 220V to analyze Cry content from total IBaCC protein and resolved Cry proteins by LC-MS, respectively. When the 50ug protein sample entered the gel, electrophoresis was stopped (to isolate unresolved total protein), while the gel with 100ug protein was fully resolved (to isolate 131-140 kDa Cry proteins). Total protein and Cry proteins were excised as stained bands and submitted to the UMass Chan Mass Spectrometry Facility (Worcester, MA) for LC-MS/MS to analyze Cry21Aa:Cry5Ba ratios in HY-BT218 expression cultures. All MS/MS samples were analyzed using Mascot v 2.1.1.21 (Matrix Science, London, UK). Scaffold v 5.3.3 (Proteome Software Inc., Portland, OR) was used to validate MS/MS based peptide and protein identifications.

### Animals and parasites

*Ancylostoma ceylanicum* hookworms were maintained in hamsters as previously described (Hu et al. 2018). Hamsters were provided with food and water ad libitum. All animal experiments with hookworms were carried out under protocols approved by the University of Massachusetts Chan Medical School (IACUC, PROTO202000071). The care and housing of animals used in this study conformed to the NIH Guide for the Care and Use of Laboratory Animals in Research (18-F22) and all requirements issued by the USDA, including regulations implementing the Animal Welfare Act (P.L. 89-544) as amended (18-F23). The procedures for fecal sample collection from sheep were approved by the Institutional Animal Care and Use Committee at the University of Rhode Island under protocol AN2021-013.

### *Ex vivo* experiments

Adult *A. ceylanicum* hookworms harvested 18 days post-inoculation (PI) from the intestines of infected hamsters were assayed *ex vivo* after 24h incubation with IBaCC treatments using the Worminator platform as previously described, with n = 8 adult hookworms/condition (Chicca et al. 2022; Hoang et al. 2024). The motility of each group was normalized to the average buffer control in hookworm medium. *Haemonchus contortus* eggs were isolated from ovine feces and used for larval development assays in triplicate (see Figure 8) as previously described (Hoang et al. 2024; Sanders et al. 2020). Results are graphed as the average of three experiments.

## Graphs and statistics

Graphs and statistics were prepared with GraphPad Prism v. 10. For Figure 8, to determine if the strains were efficacious against hookworm adults, data were analyzed using analysis of variance (ANOVA) and one-tailed Dunnett’s test using Δ*spo0A* strain as the control for comparison.

## Results

### Expression of Cry5Ba in *spo0A* and Δ*spo0A* Bt strains

We have previously shown that recombinant Cry5Ba expression can vary between *spo^+^* Bt strains (Hu et al. 2010). Therefore, we set out to determine which Bt strains might be well suited for studies with this protein. We transformed several acrystalline, *spo^+^* Bt strains (Table S1) with pHY159 (*cry5Ba*-expressing) or pHT3101 as an empty vector control (Lereclus et al. 1989) and ascertained Cry5Ba production. Relatively speaking, BMB171 had very low levels of Cry5Ba expression, 42Q had low levels of expression, and HD73-20, HD-1, and 407 each expressed Cry5Ba well (Figure S1). The detection of Cry5Ba after just 12 hours of culture demonstrates the expected vegetative *cry5Ba* gene expression from the *cry3A* promoter. We also observed more Cry5Ba in samples taken at 48h, indicating that Cry5Ba synthesis continues even after the transition to stationary phase.

We next extended our studies to non-sporulating strains, which are central to IBaCC anthelmintic development (Li et al., 2021). We took 407, HD73, and the HD1 derivative 4D8 (see Figure 1), knocked out the *spo0A* locus, and studied Cry5Ba synthesis driven by the *cry3A* promoter (Li et al., 2021). SDS-PAGE analysis of these cultured strains revealed that each of these markerless Δ*spo0A* strains harboring expression plasmids produced comparably robust levels of Cry5Ba (Figure 1). The difference between expression in HD73 and 407 seen in *spo0A*^+^ cells (Figure S1) was reduced in Δ*spo0A* cells (Figure 1). 4D8 Δ*spo0A* and HD73 Δ*spo0A* cells showed reduced Cry5Ba production compared to their congenic *spo^+^* strains and were notably lysogenic. For this reason, we moved forward with Bt 407 for further studies.

### Integration of *cry5Ba* into the *spo0A* locus of the Bt genome

Our goal is to produce a Cry-producing IBaCC (asporagenous) strain for anthelmintic therapy that does not require antibiotics for plasmid maintenance. The most efficient means to accomplish this is to perform markerless integration of the *cry* gene, *e.g.*, *cry5Ba*, into the *spo0A* locus. We thus integrated the expression cassette, the *cry5Ba* gene with promoter and 3’UTR, into the *spo0A* locus of Bt 407 using the technique described (Materials and Methods; Table S3). This Δ*spo0A*::*cry5Ba* Bt 407 strain was observed by phase-contrast microscopy, confirming the *spo*^-^ phenotype, *i.e.*, no spores, along with the presence of Cry5Ba cytosolic crystals (compare Figures 2A,B). PCR analyses of isolates confirmed the replacement of *spo0A* and with Δ*spo0A*::*cry5Ba*.

SDS-PAGE analysis of this new 407 Δ*spo0A*::*cry5Ba* strain demonstrated full-length Cry5Ba expression (Figure 3A). Using incubation conditions optimized for our plasmid based expression (Hoang et al. 2024), the 140 kDa Cry5Ba protein was detected in the integrated Δ*spo0A*::*cry5Ba* strain albeit at a lower level than the same culture volume of the Δ*spo0A*::*kan* 407 strain with plasmid-based Cry5Ba expression (Figure 3A, compare Δ*spo0A + pcry5Ba* to Δ*spo0A*::*cry5Ba*).

**Figure 3.**
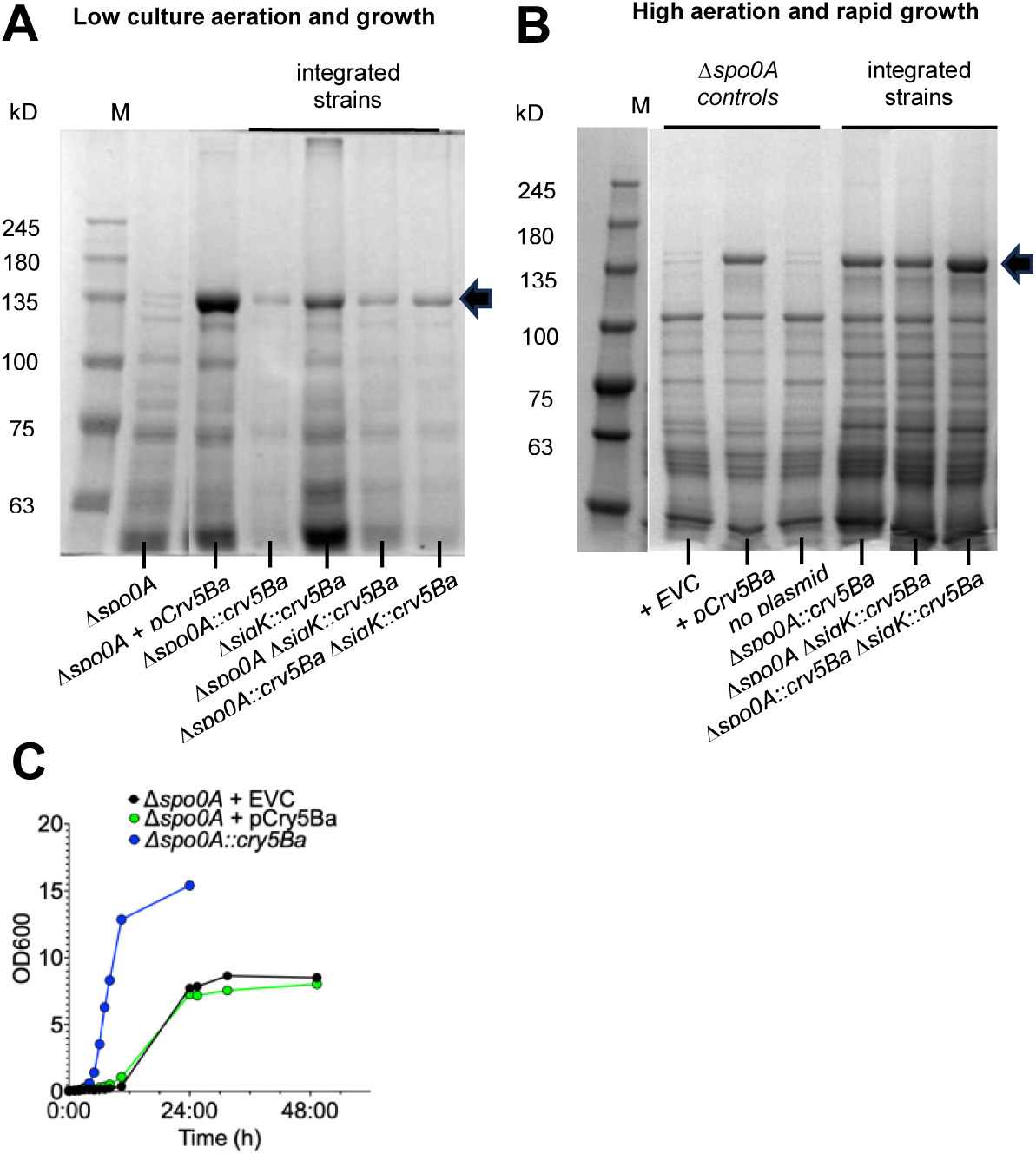
SDS-PAGE analysis of Cry-expressing strains of Bt 407. (A) Full-length Cry5Ba (arrow) in expression cultures of several Bt 407 Δ*spo0A* strains after growth with low aeration at 25 °C. Δ*spo0A*, Δ*spo0A* + p*Cry5Ba* plasmid (pHY159), Δ*spo0A*::*cry5Ba*, *ΔsigK*::*cry5Ba*, Δ*spo0A ΔsigK*::*cry5Ba,* Δ*spo0A*::*cry5Ba* Δ*sigK*::*cry5Ba*. Each lane depicts equal sample volumes (∼38ul) of 10ml in culture tubes. (B) Full-length Cry5Ba (arrow) in expression cultures of Δ*spo0A* strains after rapid growth in optimal conditions at 30 °C. Δ*spo0A* + pHT3101 (EVC), Δ*spo0A* + pCry5Ba, Δ*spo0A*, Δ*spo0A*::*cry5Ba*, Δ*spo0A ΔsigK*::*cry5Ba*, Δ*spo0A*::*cry5b* Δ*sigK*::*cry5Ba*. Each lane depicts equal sample volumes (20ul) from 25ml cultures in beveled flasks. (C) Representative growth curve during 30 °C incubations of Δ*spo0A*::*cry5Ba* (green circles) compared to plasmid-harboring strains, Δ*spo0A* + EVC (black) and Δ*spo0A* + pCry5Ba (green circles).

### Integration of *cry5Ba* into the *sigK* locus of the Bt genome

Next, we used markerless integration to insert P*_cry3A_-cry5Ba* into the *sigK* locus.

*sigK* encodes σ^K^, the last essential sigma factor involved in sporulation (Bravo et al. 1996; Hilbert and Piggot 2004; Lereclus et al. 2000). In a Δ*spo0A* background, the sporulation-dependent *sigK* is functionally moot, not expressed. We produced a pHY304 derivative with P*_cry3A_-cry5Ba* inserted into the *sigK* locus (Δ*sigK*::*cry5Ba*), designated pHY313 (see Table S3). We also produced plasmids with integrated P*_cry3A_-cry5Ba* into the *sigK* locus and then integrated Δ*sigK*::*cry5Ba* into both asporagenous Δ*spo0A* and Δ*spo0A*::*cry5Ba* cells, generating an alternate single integration and a double integrant, respectively (Table S3). The resultant new Δ*spo0A* Δ*sigK*::*cry5Ba* and Δ*spo0A*::*cry5Ba* Δ*sigK*::*cry5Ba* strains presented with uniform Cry5Ba inclusions visualized in almost all cells (Figure 2C, D). The strain with two *cry5Ba* copies presented with visibly larger crystals than strains with one *cry5Ba* copy (Figure 2B,C). No inclusion bodies are seen in the Δ*spo0A* controls (Figure 2A).

Initial expression experiments incubated in culture tubes showed that all *cry5Ba*-integrated strains produced Cry5Ba, albeit less per unit volume than our plasmid-harboring Δ*spo0A* strain. (Figure 3A). Notably, no obvious difference in Cry5Ba abundance was observed between the two Δ*spo0A* strains with a single *cry5Ba* copy at either the Δ*spo0A* or Δ*sigK locus*. However, Δ*sigK::cry5Ba* cultures exhibited more biomass and Cry5Ba content per culture volume than either Δ*spo0A::cry5Ba* or Δ*spo0A* Δ*sigK::cry5Ba*. Relative to background, slightly more Cry5Ba production could be seen in the double than the single integrant (Figure 3B).

In our efforts to increase expression from our Δ*spo0A*::*cry5Ba* strain, we hypothesized that higher agitation in baffled flasks (better aeration) and higher temperatures, for optimal Bt growth, would improve expression of chromosomally integrated *cry* genes (Figure 3B). Under these growth conditions, either integrated strain with a single copy of *cry5Ba,* Δ*spo0A*::*cry5Ba* or Δ*spo0A* Δ*sigK*::*cry5Ba,* achieved Cry5Ba abundance comparable to the plasmid-based expression strain in terms of production per unit volume of culture (Figure 3B). The integrated strain with two *cry5Ba* copies, Δ*spo0A*::*cry5Ba* Δ*sigK*::*cry5Ba,* exhibited higher Cry5Ba abundance than either strain with a single *cry5Ba* copy, Δ*spo0A*::*cry5Ba* and Δ*spo0A* Δ*sigK*::*cry5Ba*, in both culture conditions (Figures 3A, B). We quantitated the abundance of Cry5Ba protein from the single and double integrated strains grown under identical conditions. The double-*cry5Ba* integrant strain produced ∼1.9 ± 0.1 (standard deviation of the mean; n=3) times the amount of Cry5Ba protein than Δ*spo0A*::*cry5Ba* per volume of culture grown at 30 °C with high aeration.

We also compared the growth rates and maximal culture densities of the integrated and plasmid-bearing strains during culture (Figure 3C). We found that integrated Cry5Ba (Δ*spo0A*) in the absence of antibiotic achieved higher culture densities than plasmid (empty vector or *cry5Ba* insert)-bearing strains (Δ*spo0A*) with antibiotics or the isogenic Δ*spo0A* strain without any plasmid and no antibiotics.

Because the yields are similar between the strains, these results suggest that the yield per cell in the integrated strain is lower than in the plasmid-strain and provides insight into an area for increased productivity.

### Integration of a second Bt cry gene into the genome

Because Cry protein combinations are common in nature and are a way to increase efficacy, broaden tropism, and prevent resistance (Lee et al. 1996; Tabashnik, Brevault, and Carriere 2013; Schnepf et al. 1998; van Frankenhuyzen 2013; Bravo et al. 1998), we chose to integrate a second nematicidal Cry protein, Cry21Aa (Wei et al., 2004; Hu et al., 2010), alone and also in combination with Cry5Ba. Using the same allelic exchange method, we generated two *spo*^-^ *cry21Aa-*integrated strains, Δ*spo0A*::*cry21Aa* and Δ*spo0A*::*cry21Aa* Δ*sigK*::*cry5Ba* (see Table S4). We observed that Δ*spo0A*::*cry21Aa* cells do not present discernible crystals of Cry21Aa. Single crystalline inclusion bodies can be seen in the bacilli of Δ*spo0A*::*cry21Aa* Δ*sigK*::*cry5Ba* cells (Figure 2E). In both cases, only asporagenous cells were observed, no spores.

SDS-PAGE was used to analyze the protein content of these cells. Like the double integration strain with two copies of *cry5Ba,* this Δ*spo0A::cry21Aa* Δ*sigK::cry5Ba* strain expresses more total Cry protein than either Δ*spo0A* strain with a single *cry5Ba* copy (Figures 3B and 4). Full-length Cry21Aa is ∼131 kDa, slightly smaller than the ∼140kDa Cry5Ba, which could be clearly seen in strains Δ*spo0A*::*cry21Aa* and Δ*spo0A*::*cry21Aa* Δ*sigK*::*cry5Ba* (Figure 4). Densitometry of stained gel images in which the two Cry proteins are resolved suggested that the ratio of full-length Cry5Ba::Cry21Aa in Δ*spo0A*::*cry21Aa* Δ*sigK*::*cry5Ba* is approximately 2.5::1. LC-MS/MS spectral counts of both total protein samples and excised bands of Cry21Aa and Cry5Ba from the strain *spo0A*::*cry21Aa* Δ*sigK*::*cry5Ba* (see Materials and Methods) revealed ratios of full-length Cry5Ba:Cry21Aa as 2.3 and 2.7, respectively, confirming our calculations based on densitometry.

**Figure 4.**
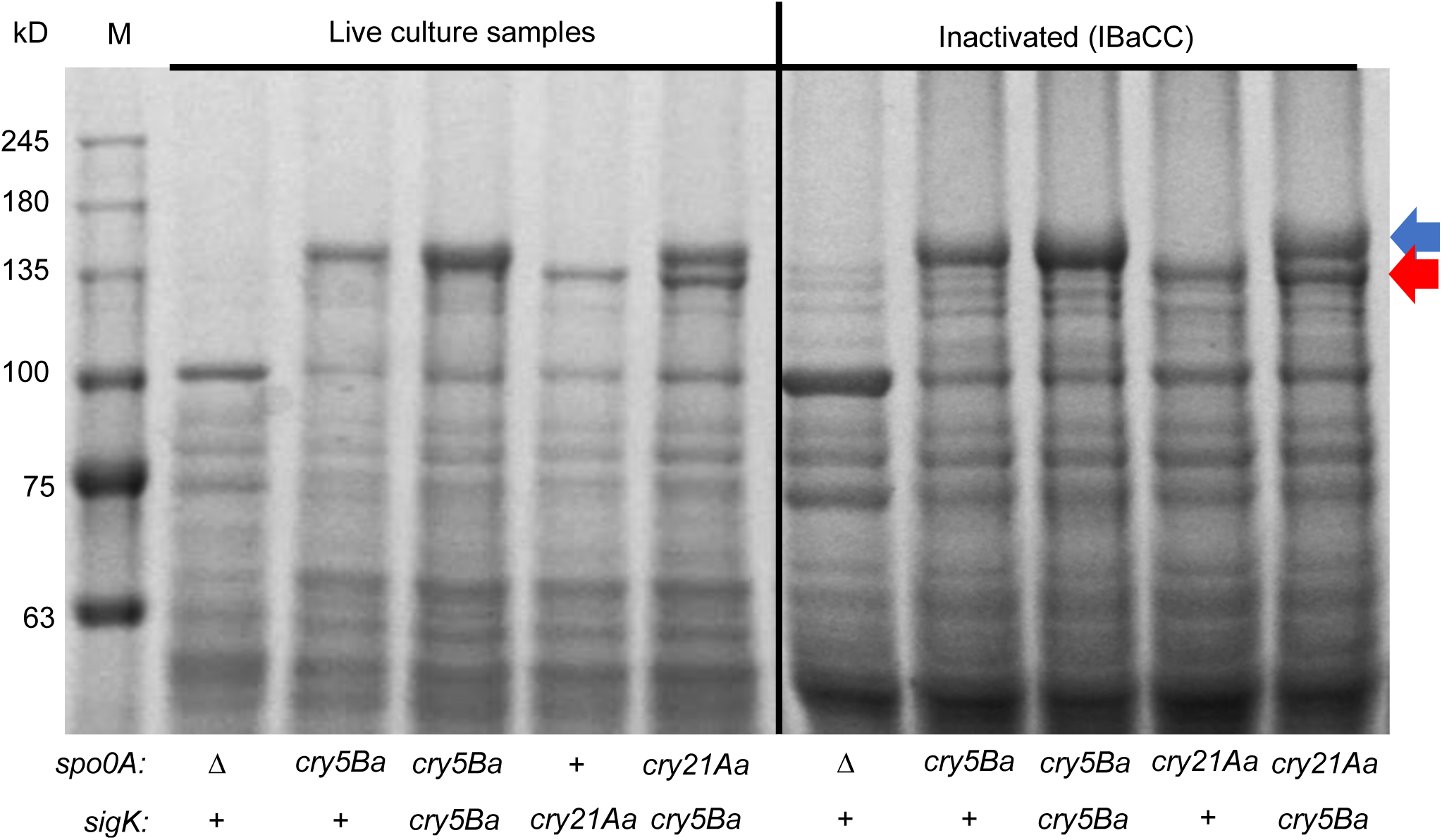
SDS-PAGE of single- and dual Cry-expressing Δ*spo0A* strains, live (left side) and inactivated (IBaCC; right side) cells. Full-length Cry5Ba (blue arrow) and Cry21Aa (red arrow). M = protein standard, Δ*spo0A*, Δ*spo0A*::*cry5Ba*, Δ*spo0A*::cry5Ba Δ*sigK*::*cry5Ba*, Δ*spo0A*::*cry21Aa*, Δ*spo0A*::*cry21Aa* Δ*sigK*::*cry5Ba*. Loading was based on equal volume of sample from equal volume of culture, or ∼38ul of 50ml flask cultures.

### Stability of integrants

We next sought to determine the stability of our double integrated strain. Our double *cry5Ba* gene insertions at the *spo0A* and *sigK* loci are approximately 145 kb apart in the approximately 5.5 Mb chromosome of Bt 407^-^(Sheppard et al. 2013), which could possibly lead to homologous recombination events and/or gene instability. To ascertain the likelihood of unwanted genomic events (i.e. recombination, deletion, mutation) at either Cry gene insertion site, we started expression cultures of each double integration strain, Δ*spo0A*::*cry5Ba* Δ*sigK*::*cry5Ba* and Δ*spo0A*::*cry21Aa* Δ*sigK*::*cry5Ba,* alongside relevant control strains. We used shake flasks for rapid growth. After an initial overnight culture, cultures were twice daily (morning after an overnight and then ∼7-8 hr later) subjected to 1000-fold dilutions into fresh media flasks to maintain logarithmic growth, for a total of six inoculations (Figure 5A). Our negative control strain, Δ*spo0A,* and each of our double integration strains, Δ*spo0A*::*cry5Ba* Δ*sigK*::*cry5Ba* and Δ*spo0A*::*cry21Aa* Δ*sigK*::Cry5Ba, exhibited similar doubling times approximating 48 minutes, (Table S5). Thus, our experiment resulted in more generations of bacterial division than the number of generations needed to fill a 100 L fermenter.

**Figure 5.**
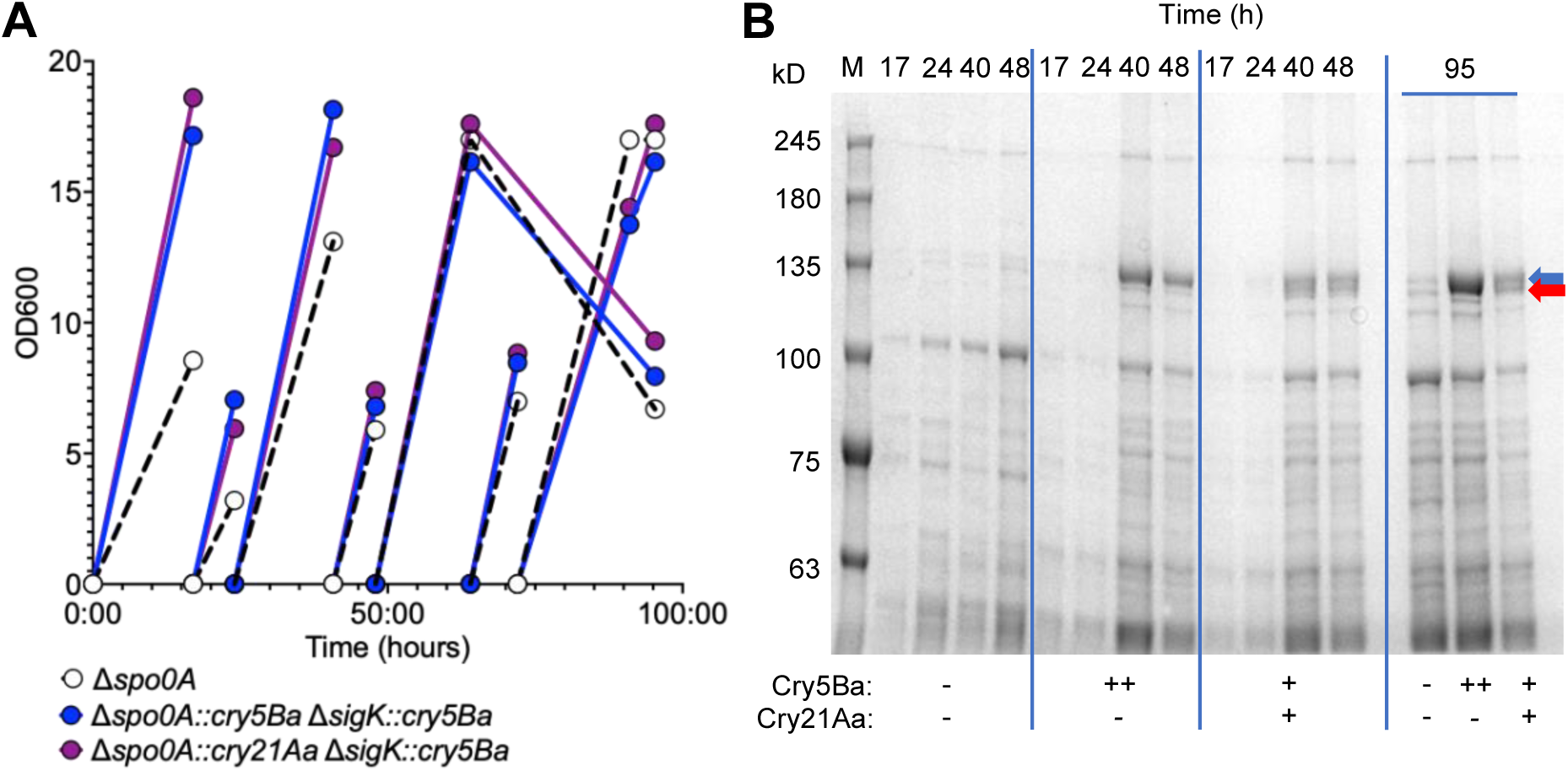
Time course of expression cultures. (A) Representative growth curves of select integrated strains, Δ*spo0A* (black line), Δ*spo0A*::*cry5Ba* Δ*sigK*::*cry5Ba* (blue), and Δ*spo0A*::*cry21Aa* Δ*sigK*::*cry5Ba* (purple). Samples from first culture flask of each strain removed for SDS-PAGE at time points 17h, 24h, 40h, and 48h. Stationary phase samples from fifth flask dilution of each strain removed at 95h time point ∼48h after inoculation (up black arrow). (B) Left three panels: SDS-PAGE analyses of Δ*spo0A*, Δ*spo0A*::*cry5b* Δ*sigK*::*cry5b,* and Δ*spo0A*::*cry21A* Δ*sigK*::*cry5B* over the first 48h of continual growth from the first inoculum. Right panel: expression found in the 95-hour samples (up arrow in 6A). Full-length Cry5B (blue arrow) and Cry21A (red arrow).

To follow expression over time, mid-log samples were also removed for SDS-PAGE analysis over the course of 48 hr (Figure 5B, left three panels) and again at the 95h end. To check stability, final samples were taken at 95 hr from the beginning of the experiment, an accumulation of >100 generation times over the course of the experiment. These final culture samples were analyzed for both Cry protein content and genomic sequences of integrated expression cassettes. Protein gel showed appropriate expression of each Cry protein as expected (Figure 5B). The high molecular weight proteins in the Δ*spo0A* control strain are not Cry proteins, though similarly sized.

Importantly, the genomes of cells within 1ml of dense culture samples were extracted and scrutinized. The integrity of each chromosomal expression locus of both doubly integrated strain Δ*spo0A*::*cry5Ba* Δ*sigK*::*cry5Ba and* Δ*spo0A*::*cry21Aa* Δ*sigK*::*cry5Ba* was confirmed by sequencing; each were intact with no mutations.

### Integrated strains are bioactive

To determine if Cry proteins generated in these integrated strains were bioactive, we generated IBaCC preparations of each strain to bioassay against parasitic nematodes (Hoang et al. 2024; Li et al. 2021; Urban et al. 2021). As shown in Figure 4 (right side), IBaCC processing resulted in no significant degradation or loss of either or both Cry proteins. Moreover, IBaCC samples of all Cry-integrated strains retained uniform crystalline inclusions in intact cells (Figure 6), as has been seen before (Hoang et la., 2024; Li et al., 2021).

**Figure 6.**
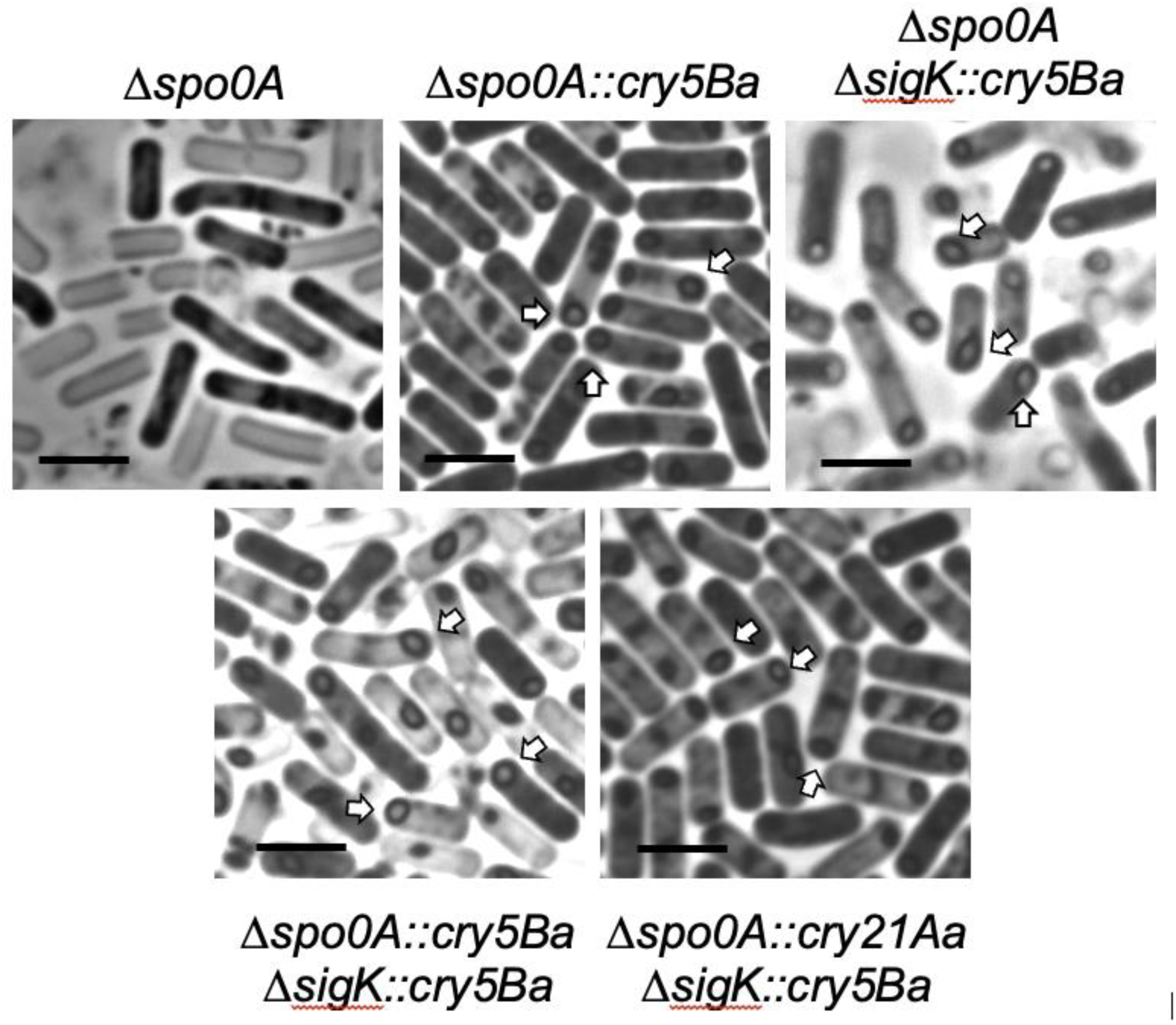
Phase-contrast microscopy of indicated markerless, asporagenous *Bt* 407 strains processed to IBaCC. Arrows point to crystal inclusions. Scale bar = 5 µm.

To ascertain bioactivity of each Cry protein initially alone, single integrants of Cry5Ba and Cry21Aa were used in dose-response larval development assay using *Haemonchus contortus* larvae, a critically important parasite of ruminants (Gilleard 2013). Cry5Ba from Δ*spo0A* Δ*sigK*::*cry5Ba* IBaCC preparations was fully bioactive, completely inhibiting iL3 development at 50 ng/ml (Figure 7), with efficacy similar to published data from a non-integrated Cry5Ba IBaCC strain (Sanders et al. 2020).

**Figure 7.**
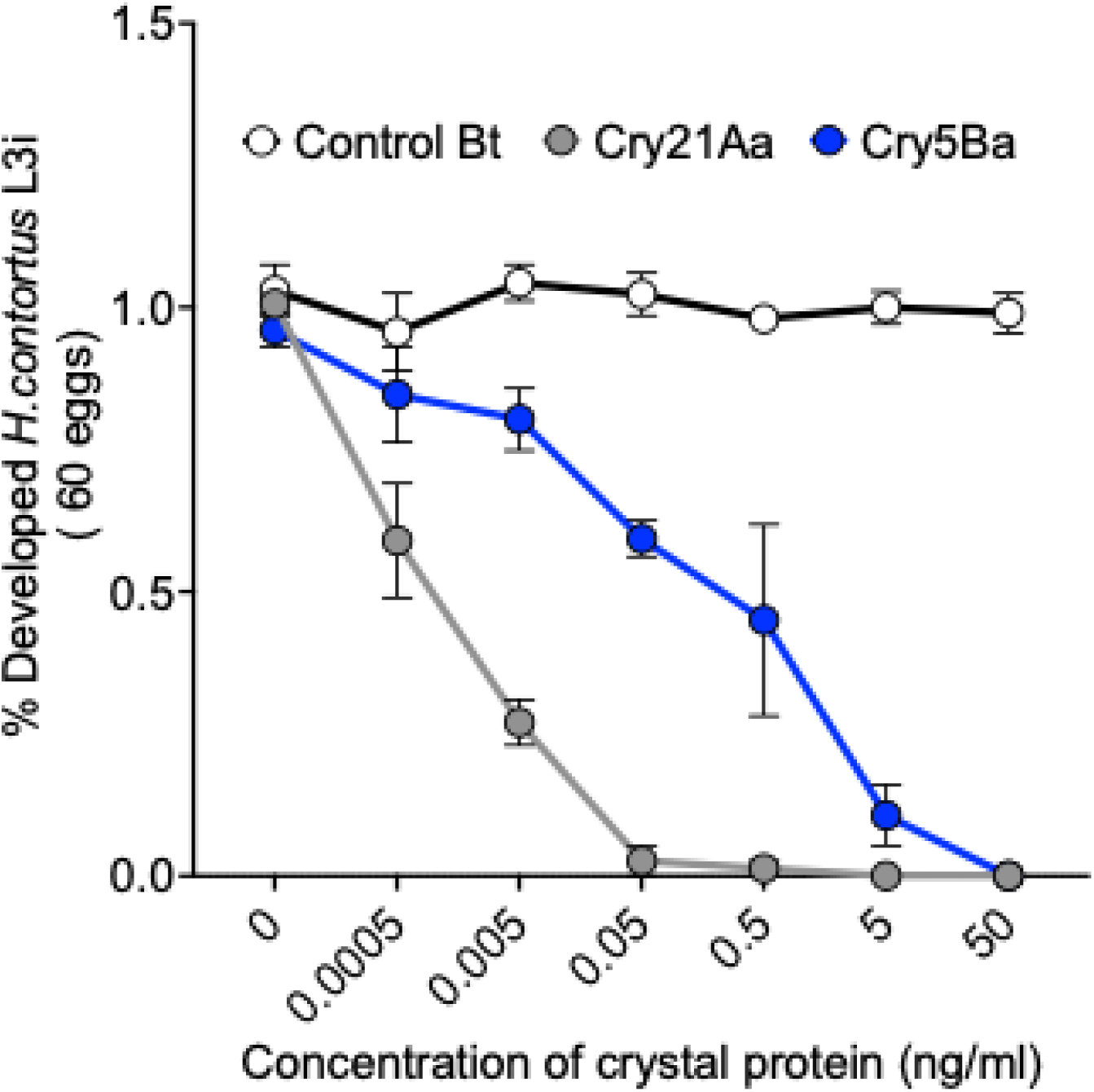
Efficacy of integrated markerless IBaCC strains against *Haemonchus contortus* larval development. Ratio of *Haemonchus contortus* eggs that develop to the infectious third larval (iL3) stage on increasing doses of proteins in IBaCC. Data are each normalized to the average of media without Bt (negative control). Control Bt = Δ*spo0A*, Cry5Ba = Δ*spo0A* Δ*sigK*::*cry5b* and Cry21Aa *=* Δ*spo0A*::*cry21A*. 40.1 ± 1.1 L3i developed from ∼ 60 eggs per well in media. n = 3 experiments, each in triplicate. Δ*spo0A* content was normalized by volume to the strain with the lowest Cry protein concentration and similarly diluted but contains no Cry protein. Plotted are the mean and standard error.

**Figure 8.**
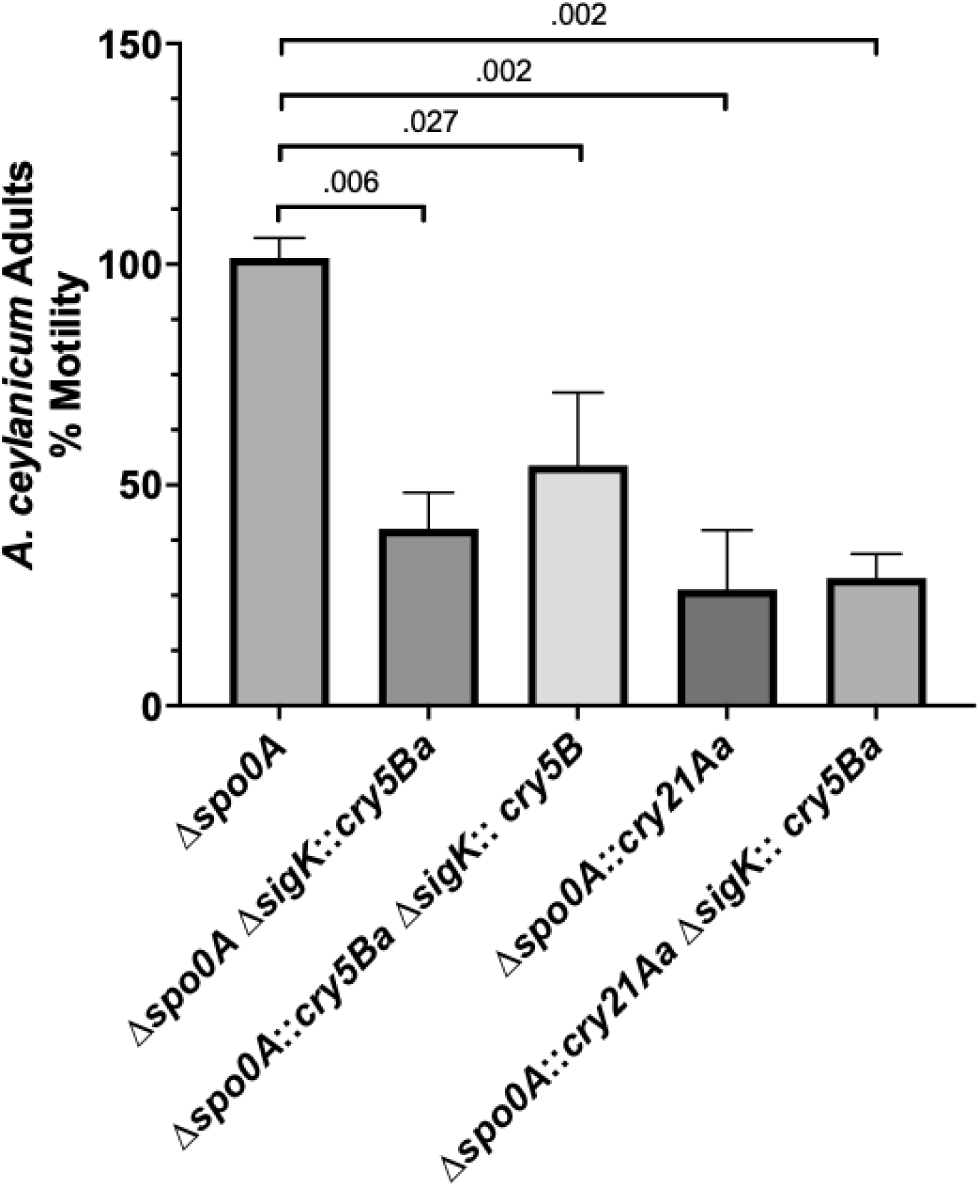
Bioactivity assays of integrated *Bt* IBaCC strains against *A. ceylanicum* hookworms *ex vivo*. Plotted is average relative hookworm motility after incubations with Cry- Δ*spo0A*, Δ*spo0A* Δ*sigK*::*cry5ba,* Δ*spo0A*::*cry5ba* Δ*sigK*::*cry5ba*, and Δ*spo0A*::*cry21Aa,* and Δ*spo0A*::*cry21Aa* Δ*sigK*::*cry5Ba* (error bars represent standard error). All motility data are normalized to the media only (amu = 117 ± 18) and represent the average of three independent experiments. The total concentration of Cry protein in each experiment was 0.5 µg/mL. Δ*spo0A* content was normalized by volume to the strain with the lowest Cry protein concentration at 0.5 ug/ml. Omitting the data for *Δspo0A*, none of the cry-integrated strains were statistically different from the single *cry5Ba* integrated strain (see Materials and Methods). Note, although the double cry5Ba integrated strain produces twice as much Cry protein as the single integrated strain, this data was collected using the same concentration of Cry5Ba protein in the assay from either strain.

Cry21Aa IBaCC preparations from the strain Δ*spo0A*::*cry21Aa* significantly inhibited iL3 development at 50 pg/mL (Figure 7). Cry21Aa was consistently 2-3 orders of magnitude more potent than Cry5Ba at inhibiting larval development, indicating that Cry21Aa is more bioactive against developing *H. contortus* larvae than Cry5Ba.

IBaCC from single and double-integrated strains were quantitated and also tested *ex vivo* against adult *A. ceylanicum* hookworms, a zoonotic hookworm parasite increasingly recognized as an important pathogen of humans (Colella, Bradbury, and Traub 2021). Relative to media control, the Cry- negative Δ*spo0A* strain had no bioactivity. In contrast, Δ*spo0A* Δ*sigK*::*cry5Ba*, Δ*spo0A*::*cry5Ba* Δ*sigK*::*cry5Ba* (double *cry5Ba* integrant), Δ*spo0A*::*cry21Aa*, and Δ*spo0A*::*cry21Aa* Δ*sigK*::*cry5Ba* (*cry21Aa* and *cry5Ba* co-integrant) all exhibited strong bioactivity against the hookworms.

### Discussion and Conclusion

We have developed an inactivated asporagenous Bt system for suitable for large scale manufacturing and safe for oral delivery of anthelmintic Cry proteins (Li et al. 2021). Since (1) this system requires the bacterium to be genetically asporagenous and (2) we want to obviate the use of antibiotic-resistance cassettes required to maintain plasmids or inserted/deleted genes—we sought to investigate markerless integration of anthelmintic Cry proteins into targeted loci of Bt.

Here we demonstrate for the first time markerless integration in Bt using methods established for other *Bacilli*. These integrants of Cry5Ba and Cry21Aa into the *spo0A* and/or *sigK* loci result in good expression of each protein as well as complete inhibition of sporulation. We also demonstrate that these proteins can be co-expressed in the same strain and that integration of two copies of the same gene, *cry5Ba*, into two different loci resulted in increased protein production.

Importantly, we were also able to demonstrate that the integrity of each P*_cry3A_*-*cry5Ba* insertion in our Δ*spo0A*::*cry5Ba* Δ*sigK*::*cry5Ba* strain remains stable well over the extended culture time required for to fill industrial-scale fermenters (>100 generations). The lack of mutations in our *cry5Ba* constructs and the sustained Cry5Ba accumulation are also indicative of stability of this integrated system. Similarly, our double Δ*spo0A*::*cry21Aa* Δ*sigK*::*cry5Ba* integrant was stable over the same number of doublings.

In addition to stable integration and expression, the Cry proteins produced in these strains are bioactive, able to intoxicate *ex vivo* both *H. contortus* parasitic larvae and parasitic hookworm adults. The efficacy of Cry5Ba from integrated strains is equivalent to that previously published. The data also indicate that Cry21Aa is also highly bioactive against both parasites, with efficacy comparable or superior to that of Cry5Ba. These are the first reports of Cry21Aa efficacy against either of these parasites.

As noted above, an advantage of our integrated system is the elimination of antibiotics. This elimination reduces production costs and avoids the introduction of residual antibiotics into the final product. Antibiotic residues can negatively affect the microbiomes of treated animals (Sadighara et al. 2023; Arsene et al. 2022; Chen et al. 2022), and the antibiotic resistance genes presented on plasmids could potentially be taken up by opportunistic pathogens (Tao et al. 2022; Jian et al. 2021).

Chromosomal integration itself offers more consistent expression per cell compared to the expression from variable plasmid numbers across cells and improved expression of proteins under conditions of rapid bacterial growth in which larger plasmids are more easily lost in the population (Grabherr et al. 2002). The absence of noncoding plasmid DNA in this system also reduces the cellular burden associated with plasmid replication, which negatively impacts growth rate and biomass accumulation (Silva, Queiroz, and Domingues 2012), a main reason that high copy number do not always imply an increase in production yields (Rosano and Ceccarelli 2014; Jones, Kim, and Keasling 2000).

A significant advantage of our methodology here is the ability to stably insert multiple Cry proteins. A large plasmid encoding multiple Cry proteins can be challenging to maintain, purify, and transform into bacteria. The alternative of maintaining multiple smaller plasmids for Cry co-expression typically requires multiple antibiotics for selection. In contrast, chromosomal integration enables the integration of multiple cry genes in a single strain, allowing for stable, low-cost Cry production without the need for large plasmids or dual antibiotic selection. Additionally, the increased protein abundance with a stepwise increase in gene copy number shown there suggest that tailoring the abundance of specific Cry proteins relative to each other could be achieved with increasing chromosomal insertions, a strategy that remains underexplored in current Bt engineering.

In summary, here we demonstrate for the first time markerless integration of Cry proteins into Bt, singly, doubly, and in combination and demonstrate its bioactivity. By utilizing sporulation loci for integration, we can take advantage of the IBaCC system for oral delivery of Cry protein anthelmintics. However, the technique could easily be used for integration into other loci. Chromosomal integration, with its independence from plasmid-based and antibiotic-based expression, offers significant advantages for Cry protein production outlined above in the expression of single or multiple Cry proteins and in the control of relative Cry protein abundance. The observed nematicidal properties of our novel Bt IBaCC strains, produced without antibiotics, further advances the low-cost, scalable, and safe production of critically needed, orally administered, new anthelmintics for the control of globally important debilitating STH and GIN parasites of humans and animals.

## Author Contributions

Conceptualization, KAF, GRO, and RVA; Methodology, KAF, NC, HL, and RVA; Validation, KF, NC, and RVA; Formal Analysis, KAF and RVA; Investigation, KAF, HL, and NC; Resources, KAF, HL, NC, LK, and KLP; Data Curation, KAF, NC and HL; Writing—Original Draft Preparation, KAF; Writing—Review & Editing, KAF, GRO, and RVA; Visualization, KAF and RVA; Supervision, KAF, GRO, and RA; Project Administration, KAF, GRO, and RVA; Funding Acquisition, RVA.

## Acknowledgements

This work was financially supported by the National Institutes of Health National Institute of Allergy and Infectious Diseases grants R01-AI056189 and R01- AI182355 to R.V.A., and USDA-NIFA-AFRI Grant no. 2021-67015-34574 from the USDA National Institute of Food and Agriculture to K.H.P. Plasmid pHY304 was kindly provided by Dr. Craig Rubens’ laboratory, Seattle Children’s Research Institute.

## References

Ahuir-Baraja, A. E., F. Cibot, L. Llobat, and M. M. Garijo. 2021. ‘Anthelmintic resistance: is a solution possible?’, Exp Parasitol, 230: 108169.

Arsene, M. M. J., A. K. L. Davares, P. I. Viktorovna, S. L. Andreevna, S. Sarra, I. Khelifi, and D. M. Sergueievna. 2022. ‘The public health issue of antibiotic residues in food and feed: Causes, consequences, and potential solutions’, Vet World, 15: 662–71.

Azizoglu, Ugur, Gholamreza Salehi Jouzani, Estibaliz Sansinenea, and Vincent Sanchis-Borja. 2023. ‘Biotechnological advances in Bacillus thuringiensis and its toxins: Recent updates’, Reviews in Environmental Science and Bio/Technology, 22: 319–48.

“Bacillus Genetic Stock Center Catalog of Strains.” In. 1999. 28. Columbus, OH.

Bravo, A., H. Agaisse, S. Salamitou, and D. Lereclus. 1996. ‘Analysis of cryIAa expression in sigE and sigK mutants of Bacillus thuringiensis’, Mol Gen Genet, 250: 734–41.

Bravo, A., S. Sarabia, L. Lopez, H. Ontiveros, C. Abarca, A. Ortiz, M. Ortiz, L. Lina, F. J. Villalobos, G. Pena, M. E. Nunez-Valdez, M. Soberon, and R. Quintero. 1998. ‘Characterization of cry genes in a Mexican Bacillus thuringiensis strain collection’, Appl Environ Microbiol, 64: 4965–72.

Cappello, M., R. D. Bungiro, L. M. Harrison, L. J. Bischof, J. S. Griffitts, B. D. Barrows, and R. V. Aroian. 2006. ‘A purified Bacillus thuringiensis crystal protein with therapeutic activity against the hookworm parasite Ancylostoma ceylanicum’, Proc Natl Acad Sci U S A, 103: 15154–9.

Chen, R. A., W. K. Wu, S. Panyod, P. Y. Liu, H. L. Chuang, Y. H. Chen, Q. Lyu, H. C. Hsu, T. L. Lin, T. D. Shen, Y. T. Yang, H. B. Zou, H. S. Huang, Y. E. Lin, C. C. Chen, C. T. Ho, H. C. Lai, M. S. Wu, C. C. Hsu, and L. Y. Sheen. 2022. ‘Dietary Exposure to Antibiotic Residues Facilitates Metabolic Disorder by Altering the Gut Microbiota and Bile Acid Composition’, mSystems, 7: e0017222.

Chicca, J., N. R. Cazeault, F. Rus, A. Abraham, C. Garceau, H. Li, S. M. Atwa, K. Flanagan, E. R. Soto, M. S. Morrison, D. Gazzola, Y. Hu, D. R. Liu, M. K. Nielsen, J. F. Urban, Jr., G. R. Ostroff, and R. V. Aroian. 2022. ‘Efficient and Scalable Process to Produce Novel and Highly Bioactive Purified Cytosolic Crystals from Bacillus thuringiensis’, Microbiol Spectr, 10: e0235622.

Colella, V., R. Bradbury, and R. Traub. 2021. ‘Ancylostoma ceylanicum’, Trends Parasitol, 37: 844–45.

Corsi, I. D., S. Dutta, A. van Hoof, and T. M. Koehler. 2020. ‘AtxA-Controlled Small RNAs of Bacillus anthracis Virulence Plasmid pXO1 Regulate Gene Expression in trans’, Front Microbiol, 11: 610036.

Crickmore, N., C. Berry, S. Panneerselvam, R. Mishra, T. R. Connor, and B. C. Bonning. 2021. ‘A structure-based nomenclature for Bacillus thuringiensis and other bacteria-derived pesticidal proteins’, J Invertebr Pathol, 186: 107438.

Deng, C., Q. Peng, F. Song, and D. Lereclus. 2014. ‘Regulation of cry gene expression in Bacillus thuringiensis’, Toxins (Basel), 6: 2194–209.

Edelstein, A. D., M. A. Tsuchida, N. Amodaj, H. Pinkard, R. D. Vale, and N. Stuurman. 2014. ‘Advanced methods of microscope control using muManager software’, J Biol Methods, 1.

Gassmann, A. J., and D. D. Reisig. 2023. ‘Management of Insect Pests with Bt Crops in the United States’, Annu Rev Entomol, 68: 31–49.

Gilleard, J. S. 2013. ‘Haemonchus contortus as a paradigm and model to study anthelmintic drug resistance’, Parasitology, 140: 1506–22.

Grabherr, R., E. Nilsson, G. Striedner, and K. Bayer. 2002. ‘Stabilizing plasmid copy number to improve recombinant protein production’, Biotechnol Bioeng, 77: 142–7.

Hilbert, D. W., and P. J. Piggot. 2004. ‘Compartmentalization of gene expression during Bacillus subtilis spore formation’, Microbiol Mol Biol Rev, 68: 234–62.

Hoang, D., K. Flanagan, Q. Ding, N. R. Cazeault, H. Li, S. Diaz-Valerio, F. Rus, E. A. Darfour, E. Kass, K. H. Petersson, M. K. Nielsen, H. Liesegang, G. R. Ostroff, and R. V. Aroian. 2024. ‘Bacillus thuringiensis Cry14A family proteins as novel anthelmintics against gastrointestinal nematode parasites’, PLoS Negl Trop Dis, 18: e0012611.

Hu, Y., S. B. Georghiou, A. J. Kelleher, and R. V. Aroian. 2010. ‘Bacillus thuringiensis Cry5B protein is highly efficacious as a single-dose therapy against an intestinal roundworm infection in mice’, PLoS Negl Trop Dis, 4: e614.

Hu, Y., T. T. Nguyen, A. C. Y. Lee, J. F. Urban, Jr., M. M. Miller, B. Zhan, D. J. Koch, J. B. Noon, A. Abraham, R. T. Fujiwara, D. D. Bowman, G. R. Ostroff, and R. V. Aroian. 2018. ‘Bacillus thuringiensis Cry5B protein as a new pan-hookworm cure’, Int J Parasitol Drugs Drug Resist, 8: 287–94.

Hu, Y., B. Zhan, B. Keegan, Y. Y. Yiu, M. M. Miller, K. Jones, and R. V. Aroian. 2012. ‘Mechanistic and single-dose in vivo therapeutic studies of Cry5B anthelmintic action against hookworms’, PLoS Negl Trop Dis, 6: e1900.

Jian, Z., L. Zeng, T. Xu, S. Sun, S. Yan, L. Yang, Y. Huang, J. Jia, and T. Dou. 2021. ‘Antibiotic resistance genes in bacteria: Occurrence, spread, and control’, J Basic Microbiol, 61: 1049–70.

Jones, A. L., R. H. Needham, and C. E. Rubens. 2003. ‘The Delta subunit of RNA polymerase is required for virulence of Streptococcus agalactiae’, Infect Immun, 71: 4011–7.

Jones, K. L., S. W. Kim, and J. D. Keasling. 2000. ‘Low-copy plasmids can perform as well as or better than high-copy plasmids for metabolic engineering of bacteria’, Metab Eng, 2: 328–38.

Jones, Kegan Romelle, and Gary Wayne Garcia. 2019. ‘Endoparasites of Domesticated Animals That Originated in the Neo-Tropics (New World Tropics)’, Veterinary Sciences, 6: 24.

Khurana, Sumeeta, Shreya Singh, and Abhishek Mewara. 2021. ‘Diagnostic Techniques for Soil-Transmitted Helminths – Recent Advances’, Research and Reports in Tropical Medicine, 12: 181–96.

Lee, M. K., A. Curtiss, E. Alcantara, and D. H. Dean. 1996. ‘Synergistic effect of the Bacillus thuringiensis toxins CryIAa and CryIAc on the gypsy moth, Lymantria dispar’, Appl Environ Microbiol, 62: 583–6.

Lereclus, D., H. Agaisse, C. Grandvalet, S. Salamitou, and M. Gominet. 2000. ‘Regulation of toxin and virulence gene transcription in Bacillus thuringiensis’, Int J Med Microbiol, 290: 295–9.

Lereclus, D., O. Arantes, J. Chaufaux, and M. Lecadet. 1989. ‘Transformation and expression of a cloned delta-endotoxin gene in Bacillus thuringiensis’, FEMS Microbiol Lett, 51: 211–7.

Li, H., A. Abraham, D. Gazzola, Y. Hu, G. Beamer, K. Flanagan, E. Soto, F. Rus, Z. Mirza, A. Draper, S. Vakalapudi, C. Stockman, P. Bain, J. F. Urban, Jr., G. R. Ostroff, and R. V. Aroian. 2021. ‘Recombinant Paraprobiotics as a New Paradigm for Treating Gastrointestinal Nematode Parasites of Humans’, Antimicrob Agents Chemother, 65.

Macaluso, A., and A. M. Mettus. 1991. ‘Efficient transformation of Bacillus thuringiensis requires nonmethylated plasmid DNA’, J Bacteriol, 173: 1353–6.

Marroquin, Lisa D, Dino Elyassnia, Joel S Griffitts, Jerald S Feitelson, and Raffi V Aroian. 2000. ‘Bacillus thuringiensis (Bt) Toxin Susceptibility and Isolation of Resistance Mutants in the Nematode Caenorhabditis elegans’, Genetics, 155: 1693–99.

Mignon, C., R. Sodoyer, and B. Werle. 2015. ‘Antibiotic-free selection in biotherapeutics: now and forever’, Pathogens, 4: 157–81.

Nester, Eugene W., Linda S. Thomashow, Matthew Metz, and Milton Gordon. 2020. “100 Years of Bacillus thuringiensis: A Critical Scientific Assessment: This report is based on a colloquium, “100 Years of Bacillis thuringiensis, a Paradigm for Producing Transgenic Organisms: A Critical Scientific Assessment,” sponsored by the American Academy of Microbiology and held November 16–18, in Ithaca, New York.” In.: American Society for Microbiology, Washington (DC).

Oliveira-Filho, E. C., and C. K. Grisolia. 2022. ‘The Ecotoxicology of Microbial Insecticides and Their Toxins in Genetically Modified Crops: An Overview’, Int J Environ Res Public Health, 19.

Pflughoeft, K. J., P. Sumby, and T. M. Koehler. 2011. ‘Bacillus anthracis sin locus and regulation of secreted proteases’, J Bacteriol, 193: 631–9.

Rakesh, V., V. K. Kalia, and A. Ghosh. 2023. ‘Diversity of transgenes in sustainable management of insect pests’, Transgenic Res, 32: 351–81.

Raymond, B., and B. A. Federici. 2017. ‘In defense of Bacillus thuringiensis, the safest and most successful microbial insecticide available to humanity - a response to EFSA’, FEMS Microbiol Ecol, 93.

Rosano, G. L., and E. A. Ceccarelli. 2014. ‘Recombinant protein expression in Escherichia coli: advances and challenges’, Front Microbiol, 5: 172.

Sadighara, Parisa, Shahrbano Rostami, Hamed Shafaroodi, Ali Sarshogi, Yeghaneh Mazaheri, and Melina Sadighara. 2023. ‘The effect of residual antibiotics in food on intestinal microbiota: a systematic review’, Frontiers in Sustainable Food Systems, 7.

Sanahuja, G., R. Banakar, R. M. Twyman, T. Capell, and P. Christou. 2011. ‘Bacillus thuringiensis: a century of research, development and commercial applications’, Plant Biotechnol J, 9: 283–300.

Sanders, J., Y. Xie, D. Gazzola, H. Li, A. Abraham, K. Flanagan, F. Rus, M. Miller, Y. Hu, S. Guynn, A. Draper, S. Vakalapudi, K. H. Petersson, D. Zarlenga, R. W. Li, J. F. Urban, Jr., G. R. Ostroff, A. Zajac, and R. V. Aroian. 2020. ‘A new paraprobiotic-based treatment for control of Haemonchus contortus in sheep’, Int J Parasitol Drugs Drug Resist, 14: 230–36.

Schindelin, J., I. Arganda-Carreras, E. Frise, V. Kaynig, M. Longair, T. Pietzsch, S. Preibisch, C. Rueden, S. Saalfeld, B. Schmid, J. Y. Tinevez, D. J. White, V. Hartenstein, K. Eliceiri, P. Tomancak, and A. Cardona. 2012. ‘Fiji: an open-source platform for biological-image analysis’, Nat Methods, 9: 676–82.

Schnepf, E., N. Crickmore, J. Van Rie, D. Lereclus, J. Baum, J. Feitelson, D. R. Zeigler, and D. H. Dean. 1998. ‘Bacillus thuringiensis and its pesticidal crystal proteins’, Microbiol Mol Biol Rev, 62: 775–806.

Sheppard, A. E., A. Poehlein, P. Rosenstiel, H. Liesegang, and H. Schulenburg. 2013. ‘Complete Genome Sequence of Bacillus thuringiensis Strain 407 Cry’, Genome Announc, 1.

Silva, F., J. A. Queiroz, and F. C. Domingues. 2012. ‘Evaluating metabolic stress and plasmid stability in plasmid DNA production by Escherichia coli’, Biotechnol Adv, 30: 691–708.

Tabashnik, B. E., T. Brevault, and Y. Carriere. 2013. ‘Insect resistance to Bt crops: lessons from the first billion acres’, Nat Biotechnol, 31: 510–21.

Tao, S., H. Chen, N. Li, T. Wang, and W. Liang. 2022. ‘The Spread of Antibiotic Resistance Genes In Vivo Model’, Can J Infect Dis Med Microbiol, 2022: 3348695.

Tinkler, S. H. 2020. ‘Preventive chemotherapy and anthelmintic resistance of soil-transmitted helminths - Can we learn nothing from veterinary medicine?’, One Health, 9: 100106.

Urban, J. F., Jr., Y. Hu, M. M. Miller, U. Scheib, Y. Y. Yiu, and R. V. Aroian. 2013. ‘Bacillus thuringiensis-derived Cry5B has potent anthelmintic activity against Ascaris suum’, PLoS Negl Trop Dis, 7: e2263.

Urban, J. F., Jr., M. K. Nielsen, D. Gazzola, Y. Xie, E. Beshah, Y. Hu, H. Li, F. Rus, K. Flanagan, A. Draper, S. Vakalapudi, R. W. Li, G. R. Ostroff, and R. V. Aroian. 2021. ‘An inactivated bacterium (paraprobiotic) expressing Bacillus thuringiensis Cry5B as a therapeutic for Ascaris and Parascaris spp. infections in large animals’, One Health, 12: 100241.

van Frankenhuyzen, K. 2013. ‘Cross-order and cross-phylum activity of Bacillus thuringiensis pesticidal proteins’, J Invertebr Pathol, 114: 76–85.

Vidal, L., J. Pinsach, G. Striedner, G. Caminal, and P. Ferrer. 2008. ‘Development of an antibiotic-free plasmid selection system based on glycine auxotrophy for recombinant protein overproduction in Escherichia coli’, J Biotechnol, 134: 127–36.

Wei, J. Z., K. Hale, L. Carta, E. Platzer, C. Wong, S. C. Fang, and R. V. Aroian. 2003. ‘Bacillus thuringiensis crystal proteins that target nematodes’, Proc Natl Acad Sci U S A, 100: 2760–5.

WHO. 2023. ‘Soil-transmitted helminth infections’.

Yim, Harry H., and Craig E. Rubens. 1998. ‘Site-specific homologous recombination mutagenesis in group B streptococci’, Methods in Cell Science, 20: 13–20.

