## Supplemental info for "Chromosomal markerless integration of anthelmintic Cry proteins into the *Bacillus thuringiensis* genome"

**Table S1. Acrystalline strains**

| Cry$^-$ Strains | Source |
|---|---|
| 407$^-$ | BGSC 4A12 |
| HD73-20 | BGSC 4D22 |
| HD-1 | BGSC 4D24 |
| 4Q2-81 | BGSC 4Q7 |
| BMB171 | BGSC 4XX6 |
| 4D8 | BGSC 4D8 |
| 407 Δ*spo0A*::*kan* | (Lereclus et al. 1995) |

*Kan*, kanamycin resistant; na, not applicable

**Table S2. Primers used in this study.**

| Primer sequence (5' → 3') | Description |
|---|---|
| CGAGGTCGACGGTATCGATAGGTGAATCGTCTTATCGAATTGGC | pHY304-*spo0A*-5'f |
| CGGCCGCTCTAGAACTAGTGTTTTATTGAACGTGCGGAG | pHY304-*spo0A*-3'r |
| GCGATAAACTCAGCGACAATTGTGTGAAAATTCCC | *spo0A*-5'r |
| CACAATTGTCGCTGAGTTTATCGCAATGGTTGC | *spo0A*-3'f |
| TCGAGGTCGACGGTATCGATAAGCTTAAGAGCCTTTAGTGTGACTG | pHY304-*sigK*-5'f |
| CTCTGCCCTTACAAACTCGTGAGGTTGTTTGTGTCTGTAC | *sigK*-5'r |
| GTACAGACACAAACAACCTCACGAGTTTGTAAGGGCAGAG | *sigK*-3'f |
| GCGGCCGCTCTAGAACTAGTGGATCCGAACAGTGTCTCTCTCATGC | pHY304-*sigK*-3'r |
| AACTCAGGATATCCGACAATTGTGTGAAAATTCCC | EcoRV-*spo0A*-5'r |
| ATTGTCGGATATCCTGAGTTTATCGCAATGGTTGCG | EcoRV-*spo0A*-3'f |
| AGGGAATTTTCACACAATTGTCGGATGAAACCTTAGATAAAAGTGC | Δ*spo0A*-Pcry3A-f |
| CGCAACCATTGCGATAAACTCAGTATAGGGTGCATAATAGTAAAGG | Δ*spo0A*-5Bterm-r |
| TCTAATACACTTTAGATATCGAGGTTGTTTGTGTCTGTAC | EcoRV-*sigK*-5'r |
| CCTCGATATCTAAAGTGTATTAGAAGCAGCC | EcoRV-*sigK*-3'f |
| GTACAGACACAAACAACCTCGATGAAACCTTAGATAAAAGTGC | Δ*sigK-Pcry3A*-f |
| CTGCTTCTAATACACTTTATATAGGGTGCATAATAGTAAAGG | Δ*sigK*-5Bterm r |

**Table S3. Plasmids used in this study.**

| Plasmid | Description | Source |
|---|---|---|
| pHT3101 | Empty shuttle vector, *erm amp* | (Lereclus et al. 1989) |
| pHY304 | Temperature-sensitive integration plasmid *erm* | (Jones, Needham, and Rubens 2003; Pflughoeft, Sumby, and Koehler 2011) |
| pHY159 | $P_{cry3A(-635\ to\ +18)}$-*cry5Ba erm amp* | (Li et al. 2021) |
| pHY300 | $P_{cry3A(-635\ to\ +0)}$-*cry5Ba erm amp* | This study |
| pHY301 | $P_{cry3A(-635\ to\ +0)}$-*cry21Aa erm amp* | This study |
| pHY302 | pHY304-Δ*spo0A*(Bt 407) *erm* | This study |
| pHY303 | pHY304-Δ*spo0A*(Bt HD) *erm* | This study |
| pHY305 | pHY304-Δ*sigK*(Bt 407) *erm* | This study |
| pHY306 | pBluescript_Bt 407 Δ*spo0A*::EcoRV *amp* | This study |
| pHY307 | pBluescript_Bt 407 Δ*spo0A*::*cry5Ba amp* | This study |
| pHY308 | pBluescript_Bt 407 Δ*spo0A*::*cry21Aa amp* | This study |
| pHY309 | pHY304-Δ*spo0A*::*cry5Ba erm* | This study |
| pHY310 | pHY304-Δ*spo0A*::*cry21Aa erm* | This study |
| pHY311 | pBluescript_Bt 407 Δ*sigK*::EcoRV *amp* | This study |
| pHY312 | pBluescript_Bt 407 Δ*sigK*::*cry5Ba amp* | This study |
| pHY313 | pHY304-Δ*sigK*::*cry5Ba erm* | This study |

*Erm*, erythromycin resistant; *amp*, ampicillin resistant. All numbers refer to nucleotide number relative to the start codon ATG.

**Table S4. Experimental strains generated.**

| Strain | Genotype | Recipient strain | Donor plasmid |
|---|---|---|---|
| HY-BT1 | + $P_{cry3a}$-*cry5Ba erm* | HD73-20 | pHY159 |
| HY-BT3 | + $P_{cry3a}$-*cry5Ba erm* | 407 Cry- | pHY159 |
| HY-BT5 | + $P_{cry3a}$-*cry5Ba erm* | HD-1 Cry- | pHY159 |
| HY-BT7 | + $P_{cry3a}$-*cry5Ba erm* | BMB171 | pHY159 |
| HY-BT9 | + $P_{cry3a}$-*cry5Ba erm* | 4Q2-81 | pHY159 |
| HY-BT17 | Δ*spo0A* | 407 Cry- | pHY302 |
| HY-BT19 | Δ*spo0A* | HD73-20 | pHY303 |
| HY-BT21 | + $P_{cry3a}$-*cry5Ba erm* | HD-1 4D8 | pHY159 |
| HY-BT27 | Δ*spo0A* | HD-1 4D8 | pHY303 |
| HY-BT35 | Δ*spo0A* + pHT3101 *erm* | HY-BT19 | pHT3101 |
| HY-BT36 | Δ*spo0A* + $P_{cry3a}$-*cry5Ba erm* | HY-BT19 | pHY159 |
| HY-BT38 | Δ*spo0A* + pHT3101 *erm* | HY-BT27 | pHT3101 |
| HY-BT39 | Δ*spo0A* + $P_{cry3a}$-*cry5Ba erm* | HY-BT27 | pHY159 |
| HY-BT43 | Δ*spo0A* + pHT3101 *erm* | HY-BT17 | pHT3101 |
| HY-BT44 | Δ*spo0A* + $P_{cry3a}$-*cry5Ba erm* | HY-BT17 | pHY159 |
| HY-BT90 | Δ*sigK* | 407 Cry- | pHY305 |
| HY-BT111 | Δ*spo0A* + $P_{cry3a}$-*cry5Ba erm* | HY-BT17 | pHY300 |
| HY-BT146 | Δ*spo0A*::*cry5Ba* | 407 Cry- | pHY309 |
| HY-BT162 | Δ*sigK*::*cry5Ba* | 407 Cry- | pHY313 |

| | | | |
|---|---|---|---|
| HY-BT163 | Δ*spo0A* Δ*sigK*::*cry5Ba* | HY-BT17 | pHY313 |
| HY-BT164 | Δ*spo0A*::*cry5Ba* Δ*sigK*::*cry5Ba* | HY-BT146 | pHY313 |
| HY-BT215 | Δ*spo0A*::*cry21Aa* | 407 Cry- | pHY310 |
| HY-BT218 | Δ*spo0A*::*cry21Aa* Δ*sigK*::*cry5Ba* | HY-BT162 | pHY310 |

*Erm*, erythromycin resistant

**Table S5. Growth statistics of integrated strains.**

| Strain | Genotype | Doubling time (min) ± SEM |
|---|---|---|
| HY-BT17 | Δ*spo0A* | 48.4 ± 2.1 |
| HY-BT164 | Δ*spo0A*::*cry5Ba* Δ*sigK*::*cry5Ba* | 48.0 ± 1.4 |
| HY-BT218 | Δ*spo0A*::*cry21Aa* Δ*sigK*::*cry5Ba* | 48.3 ± 1.3 |

Generation times and number were calculated using the formula D = t/n. n = number of generations, t = total time, and D = doubling time. n = $(\log N_t - \log N_0)/\log 2$, where $N_t$ = $OD_{600}$ at time t; and $N_0$ = original $OD_{600}$ at inoculation.

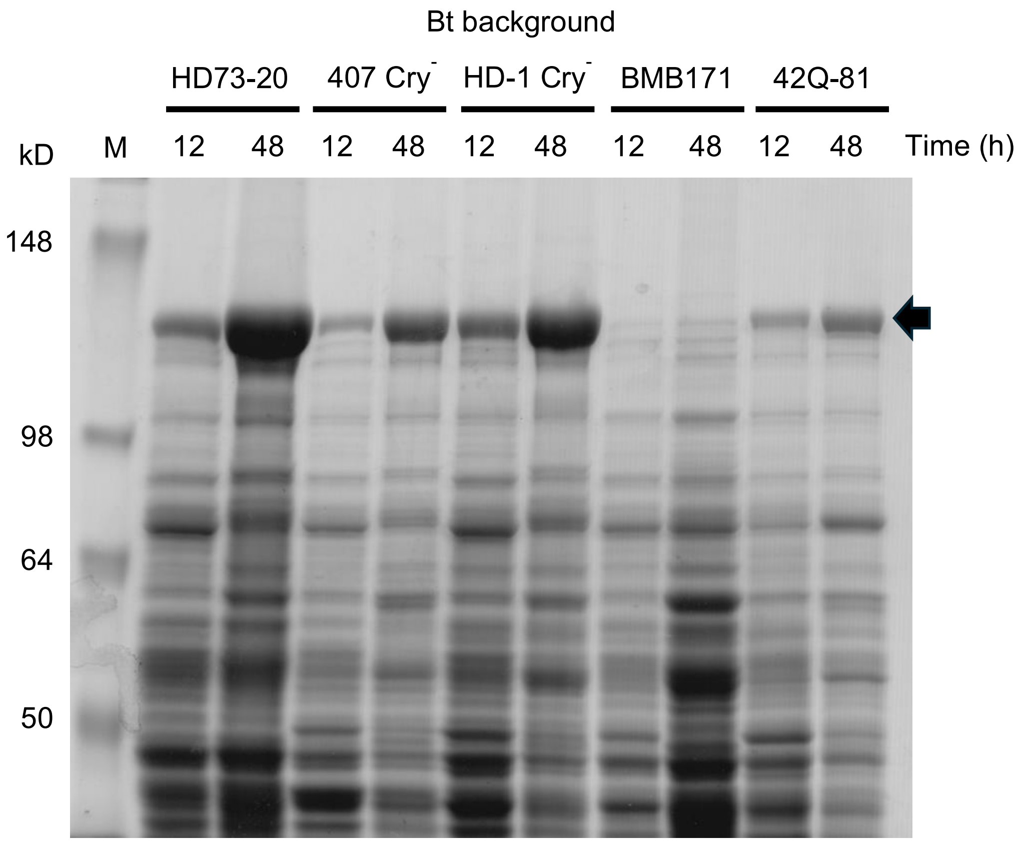


**Figure S1.** Comparison of *spo*+ *Bt* strains for Cry5B content. Cultures of several natural, acrystalline strains of *Bt* abundantly express Cry5Ba (arrow; via plasmid) to varying extent within 12h and accumulates by 48h of culture in LB. HD73 + pHY159 (strain HY-BT1), 407 + pHY159 (strain HY-BT3), HD1 + pHY159 (strain HY-BT5), BMB171 + pHY159 (strain HY-BT7), 42Q + pHY159 (strain HY-BT9).

**References**


Jones, A. L., R. H. Needham, and C. E. Rubens. 2003. 'The Delta subunit of RNA polymerase is required for virulence of Streptococcus agalactiae', *Infect Immun*, 71: 4011-7.

Lereclus, D., H. Agaisse, M. Gominet, and J. Chaufaux. 1995. 'Overproduction of encapsulated insecticidal crystal proteins in a Bacillus thuringiensis spo0A mutant', *Biotechnology (N Y)*, 13: 67-71.

Lereclus, D., O. Arantes, J. Chaufaux, and M. Lecadet. 1989. 'Transformation and expression of a cloned delta-endotoxin gene in Bacillus thuringiensis', *FEMS Microbiol Lett*, 51: 211-7.

Li, H., A. Abraham, D. Gazzola, Y. Hu, G. Beamer, K. Flanagan, E. Soto, F. Rus, Z. Mirza, A. Draper, S. Vakalapudi, C. Stockman, P. Bain, J. F. Urban, Jr., G. R. Ostroff, and R. V. Aroian. 2021. 'Recombinant Paraprobiotics as a New Paradigm for Treating Gastrointestinal Nematode Parasites of Humans', *Antimicrob Agents Chemother*, 65.

Pflughoeft, K. J., P. Sumby, and T. M. Koehler. 2011. 'Bacillus anthracis sin locus and regulation of secreted proteases', *J Bacteriol*, 193: 631-9.
